# SMOLM-LFM: ratiometric single molecule orientation without polarizers

**DOI:** 10.1101/2025.11.26.690201

**Authors:** Arturo G. Vesga, Luis Aleman-Castaneda, Hannah S. Heil, Sam Daly, Ezra Bruggeman, Steven F. Lee, Sophie Brasselet, Ricardo Henriques

## Abstract

Single-molecule orientation and localization microscopy (SMOLM) enables the determination of molecular orientation, wobbling, and position. However, most SMOLM implementations rely on complex point spread function (PSF) fitting, which limits analysis throughput and introduces high computational cost. A way to overcome these limitations is to simplify the analysis using a ratiometric intensity estimation, often relying on polarization projections. While effective in 2D, extending these methods to 3D remains challenging. Here, we introduce a new ratiometric strategy for SMOLM in 3D. Building on the principles of Single Molecule Light Field Microscopy, which captures the 3D position information from a single snapshot by segmenting the back focal plane, we extend this strategy to orientation retrieval. Our approach uses the generalized 3D Stokes formalism to linearly decompose the intensity measurements across the light-field channels, allowing computationally-efficient estimations, while avoiding both complex PSF fitting and polarization projections. This framework, called SMOLM-LFM, enables 6D estimation of single molecules (3D position + 3D orientation) with a simplified optical setup and a large depth-of-field. We present the theoretical foundations, experimental implementation, and validation through measurements on calibration beads, single fluorophores, and cells, thereby demonstrating the method’s potential and practical limitations.

## Introduction

The increasing availability of rigidly attached fluorophores has made it possible to estimate the orientation of the structures they label (1, 2). Particularly, Single Molecule Orientation Localization Microscopy (SMOLM) not only can localize individual molecules with nanometer precision, but can also determine their orientation and rotational dynamics (2–4). This enables the full structural imaging of a sample, which is key for: studying anisotropic molecular structures; understanding protein conformational changes; mapping the organization and dynamics of cellular components at the nanoscale; and providing a deeper understanding of molecular mechanisms and structural biology. Traditionally, orientational microscopy is achieved by exploiting the polarization dependence of the excitation and/or of the emission of fluorescent molecules (5), thus these techniques are generally called polarization microscopy. Since SMOLM techniques operate in a single-shot regime, they generally rely on custom detection schemes that enable evaluation of distinctive emission intensities and polarization signatures. Although many polarization-splitting schemes are limited to in-plane projections of orientation and position (6, 7), there have been recent developments to measure 3D orientations (8) or to estimate the in-depth position (9). On the other hand, point spread function (PSF) engineering techniques have been proven to estimate simultaneously both the 3D orientation and 3D position of single emitters with high precision (10, 11). However, they require more computationally intensive analysis and more complex instrumentation, and they are less robust to aberrations and polarization distortions (12, 13).

In this context, Single Molecule Light Field Microscopy (SMLFM) has been shown to be capable of estimating the 3D spatial localization of emitters over a long axial range (14–16) without dramatically increasing the PSF size, relative to comparable PSF engineering techniques (17–19). The working principle of SMLFM, which is based on Fourier-LFM (FLFM), consists in using a microlens array (MLA) at the back focal plane of the objective (BFP) to generate multiple images, each coming from different sections of the BFP. So far, SMLFM has been dedicated to the estimation of the axial position of single emitters from the estimation of the relative lateral displacement of their different sub-images. However, interestingly, the intensity distribution at the BFP is orientation-dependent (2, 20). Therefore, splitting the BFP into different sectors is also orientation-sensitive if the corresponding relative sub-images intensities of each single emitter are estimated.

In this work, we present SMOLM-LFM, a natural extension of SMLFM, a method capable of measuring the 3D orientation and 3D position of single emitters in a single snapshot without using any PSF engineering or polarization-splitting schemes. This eliminates the need for polarization-sensitive optics or phase/polarization masks, enabling a simple, inline, and compact system that can be added to most commercial microscopes as a FLFM detection module. Note that most SMOLM techniques exploit polarization and/or require polarized detection, and just a few have avoided this (3). Additionally, differently from PSF engineering techniques, which are based on a multi-parameter fit where axial position and orientation are embedded in a intricate interplay, SMOLM-LFM avoids complex and computationally costly PSF fitting since the orientation and the axial position are extracted from independent analyzes and have largely independent readouts, thus simplifying the retrieval to first approximation. We demonstrate this with simulations and experiments on three different type of samples: fluorescent nanobeads, single fluorophores deposited on a cover slip and actin labelled filaments in cells.

## Results

### Principle of SMOLM-LFM

Figure 1a shows a schematic of the experimental setup, which corresponds to that of FLFM configuration (21), similar to the previously published SMLFM works (15, 16). The key component is the MLA that is placed at a relayed plane of the BFP of the objective. Each microlens forms an individual image onto the detector, i.e., each sub-image/channel is the result of a different section of the BFP. The detection NA is set below the supercritical angle fluorescence (SAF) limit to avoid unwanted position/orientation coupling (22). As a result, each section of the BFP, which corresponds effectively to a portion of the farfield emission of the individual emitters (5), is probed in two very different ways. Firstly, the intensity distribution, which varies depending on the orientation and wobble (extent of the orientation fluctuations) of the emitter, results in distinct intensity ratios between the PSFs of the BFP sections, as depicted in Fig. 1b and c. Secondly, an axial displacement of the emitters results in a lateral displacement of the PSFs as shown in Fig. 1d, which corresponds to the common mechanism that SMLFM uses to estimate the axial position. Note that 3D position and 3D orientation estimation are theoretically uncoupled: the dipole orientation determines the intensity distribution at the BFP, while the emitter’s position affects only the phase distribution (introducing defocus and tilt in the wavefront), provided that SAF is blocked or not collected (23). Figure 1d depicts the corresponding PSFs for a fixed oblique emitter at different depths, considering a MLA made of 7 sectors, illustrating their unequal intensities among the 7 detected channels as well as their axial-dependent displacement in the image plane.

**Fig. 1.**
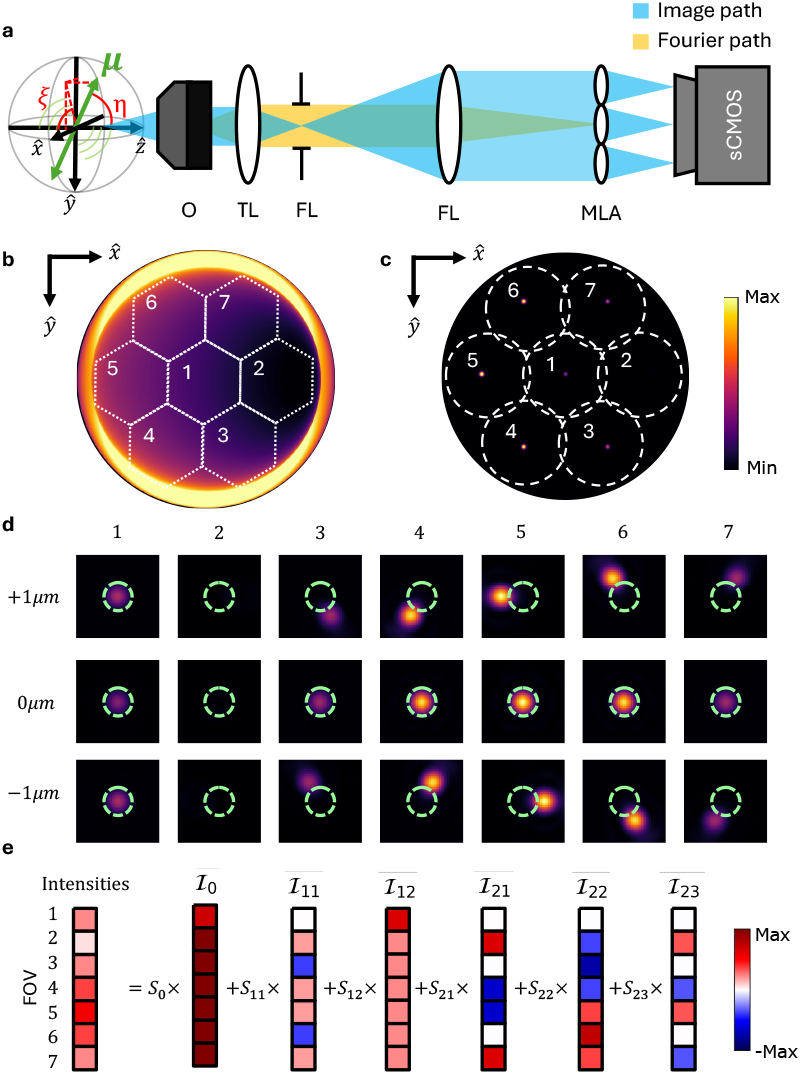
Optical setup and principle of SMOLM-LFM imaging. **a**, Sketch of the detection path of the microscope, where the emission of a fluorophore with the dipole moment ***µ*** is collected by an objective (O) that forms a first image with the tube Lens (TL). A Fourier lens (FL) relays the back focal plane (BFP) of the objective to a microlens array (MLA) that forms multiple sub-images at the detector (sCMOS). A field stop (FS) delimits the shape and size of each channel formed. Note that the dipole direction is described by the in-plane (projection on the transverse plane) and out-of-plane angles *ξ* and *η* respectively. **b**, Simulated intensity distribution at the BFP for a single dipole emitter with angles *ξ* = 0° and *η* = 45°, respectively. **c**, Image plane showing the seven perspective channel generated by the MLA. **d**, Point spread function (PSF) intensity profiles measured in each channel for the same fixed emitter but at three different in-depth positions. Note that defocus do not affect the intensity ratios but only causes lateral displacements for the external channels. In light green the respective Airy disk ≈ 1.9 µm. **e**, The measured intensities across the different channels can be linearly decomposed on the Stokes basis elements ℐ*ij* obtaining the generalized Stokes coefficients *Sij* that indicate the 3D orientation.

The measured intensity ratio, which could be obtained either from box-integration or 2D-Gaussian fitting, can be linearly decomposed using the generalized Stokes basis formalism (10, 24), which offers a fast and non-computationally heavy 3D orientation estimation. Figure 1e illustrates this decomposition, where the 7 measured channels’ intensities are translated into a linear decomposition onto known intensitybases, *I*_*ij*_, with weights being the orientation-dependent generalized Stokes parameters, *S*_*ij*_ (24). More details on the angles retrieval from these parameters are given in the Supplementary Note 1.

### Numerical simulation validation

As discussed in the literature (24, 25), only six of the nine generalized Stokes parameters are required to fully characterize 3D linear dipolar emission (orientation and wobbling). Consequently, at least six independent intensity measurements or channels are needed, a condition satisfied here through our seven-channel splitting scheme (corresponding to the MLA used). Given the linear decomposition of the measured intensities in the Stokes basis, a first and simple quantity to evaluate the performance of SMOLM-LFM is the condition number of the basis matrix (i.e. the matrix constituted by the basis elements), which is found to be *κ* ≳ 5 for a NA close-to but below SAF; refer to the Supplementary Note 1 and Fig. S1 for a detailed discussion. However, to better and further evaluate the performance of the technique, a Monte Carlo simulation is carried out reflecting empirically-determined experimental conditions, i.e., 20k photons and SNR= 7 (for more constrained conditions see Supplementary Fig. S5 and S6). For these simulations, a wobble-in-a-cone model is assumed, meaning that the orientation of the molecular dipole explores, during the integration time of the camera, a cone of aperture angle *δ* and mean orientation (*η, ξ*), as shown in Fig. 2a. To avoid the non-linearity associated with *δ*, this numerical analysis uses the degree of polarization (DoP) instead of *δ*, which, under the cone model, is given by Eq. S10. Note that within this model only four parameters are needed to describe the orientation, however we estimate the whole set of Stokes parameters to continue using a linear decomposition and avoid non-linear optimizations (see Supplementary Note 1). Due to symmetry, the simulations are limited to one quarter of the orientation sphere, since the remaining regions are equivalent. For each set of orientation parameters, 40 realizations are performed. The results shown in Fig. 2 correspond to fixed dipoles (DoP=1). For wobbling dipoles, refer to Supplementary Fig. S6.

**Fig. 2.**
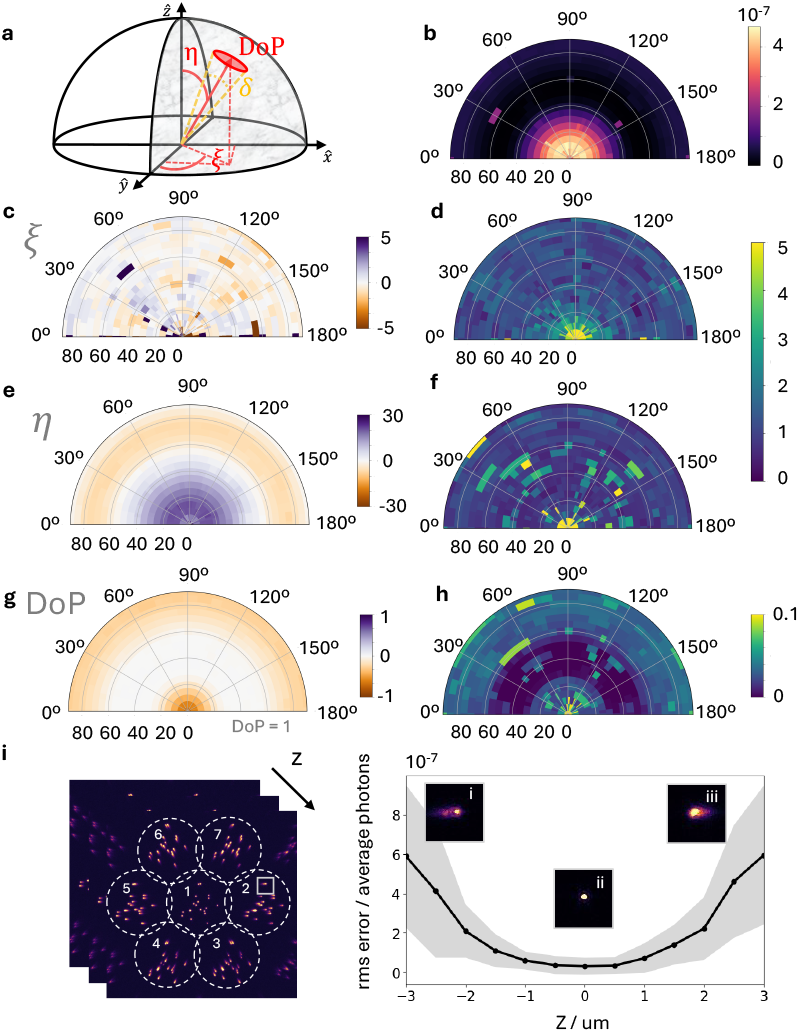
Monte Carlo simulations of fluorophore dipole orientation retrieval (20k photons, SNR=7, DoP=1). **a**, Schematic of a dipole average orientation angles: inplane angle, *ξ*; out-of-plane, *η*; and wobbling cone angle *δ*, which ultimately is related to the degree of polarization (DoP). **b**, Normalized root-mean-square (RMS) error of the intensity-ratios fitting using the Stokes-basis decomposition, normalized by the average photon count for each condition. **c–h**, Bias (left) and STD (right) of the retrieved parameters: (c,d) *ξ*; (e,f) *η*; and (g,h) DoP. **i, Left:** Representative frame of the simulated SMLFM dataset at *z* = −3 *µ*m showing individual emitters. **Right:** Corresponding normalized RMS (shaded regions indicate the standard deviation of the data). Insets show representative PSFs from channel 2 at axial positions of −3 *µ*m (i), 0 *µ*m (ii), and +3 *µ*m (iii).

Note that this Monte Carlo simulation simultaneously evaluates both the detection and orientation parameters retrieval algorithms used in the experimental setup. Figure 2b, shows the RMS error of the fitted intensities using the Stokes-basis decomposition (see Supplementary Note 1) normalized by the average number of photons per condition, and does not represent the residual error of the individual PSF fitting. As it can be seen, dipoles that are nearly out-of-plane show the largest RMS fitting error, whereas oblique orientations yield the most accurate fittings. In Figs. 2c, e and g we plot the accuracy in the estimation of the orientation parameters. While the in-plane angle, *ξ*, is estimated with a high accuracy (a few degrees), the out-of-plane angle, *η*, tends to be under/overestimated for either in-/out-of-plane dipoles (by around 10 − 20°). Part of this behaviour is expected since those are the edges of the interval in which *η* is defined. Interestingly, in these regions, for the degree of polarization, DoP, there are significant biases. On the other hand, Fig. 2d, f and h show that the standard deviation (STD) of the estimation is within several degrees of precision for most of the orientations. As expected, the in-plane angle precision decreases when dipoles lie along the longitudinal direction since, at *η* = 0°, *ξ* is not well defined. For what concerns the DoP, the STD is below 0.1 for all orientation conditions.

Lastly, we analyzed the RMS fitting error for emitters positioned at different axial positions over a 6 µm range. Individual emitters randomly oriented and with all possible wobbling states were simulated using Monte Carlo methods with the same parameters as the former simulation (20k photons, SNR= 7). The emitters were additionally positioned at different *z* positions. As shown in Fig. 2i right, the RMS fitting error increases for out-of-focus emitters. This behavior is expected, as PSFs become progressively blurred and broadened in the outer channels, reducing the accuracy of the 2D Gaussian approximation and consequently degrading both localization precision and intensity-ratio measurements, which impacts on the orientation parameters retrieval. Additional simulation conditions, including alternative MLA configurations, are provided in Supplementary Fig. S3.

### Experimental validation

An initial experimental validation and calibration of SMOLM-LFM was conducted using polarization photoselection. The probability of exciting a fluorophore is highest when the excitation polarization is parallel to the orientation of its absorption dipole, 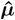 12). Controlling the illumination polarization state in an isotropic ensemble of fluorophores is therefore a possible way to select specific dipole’s orientations. Two different samples were used: fluorescent nanobeads and single fluorophores in water deposited on a coverslip (see Materials and Methods).

Fluorescent nanobeads (Tetraspeck, Thermofisher Scientific) contain densely packed fluorophores with random orientations. Energy transfer between neighboring fluorophores can occur, leading to partial depolarization of the emitted fluorescence. However, when multiple dyes or fluorophore species are present (as for tetraspeck beads), this energy transfer is hindered, and the emission remains only partially depolarized ( effectively resembling that of a highly wobbling dipole with a mean orientation aligned with the excitation polarization). This makes fluorescent beads a convenient calibration tool, as they mimic bright, wobbly dipoles in any desired direction defined solely by the illumination scheme, something that is much harder to achieve with single fluorophores whose orientations are difficult to control.

Figures 3a and b illustrate the illumination configurations and corresponding polarization states used for photoselection: two orthogonal linear polarizations at normal incidence (s in yellow and p in blue) and at near-Total Internal Reflection (TIRF) incidence (s remaining in-plane and p becoming longitudinal). The effect of each configuration is shown on tetraspeck beads and on single emitters (Fig. 3 a, b).

**Fig. 3.**
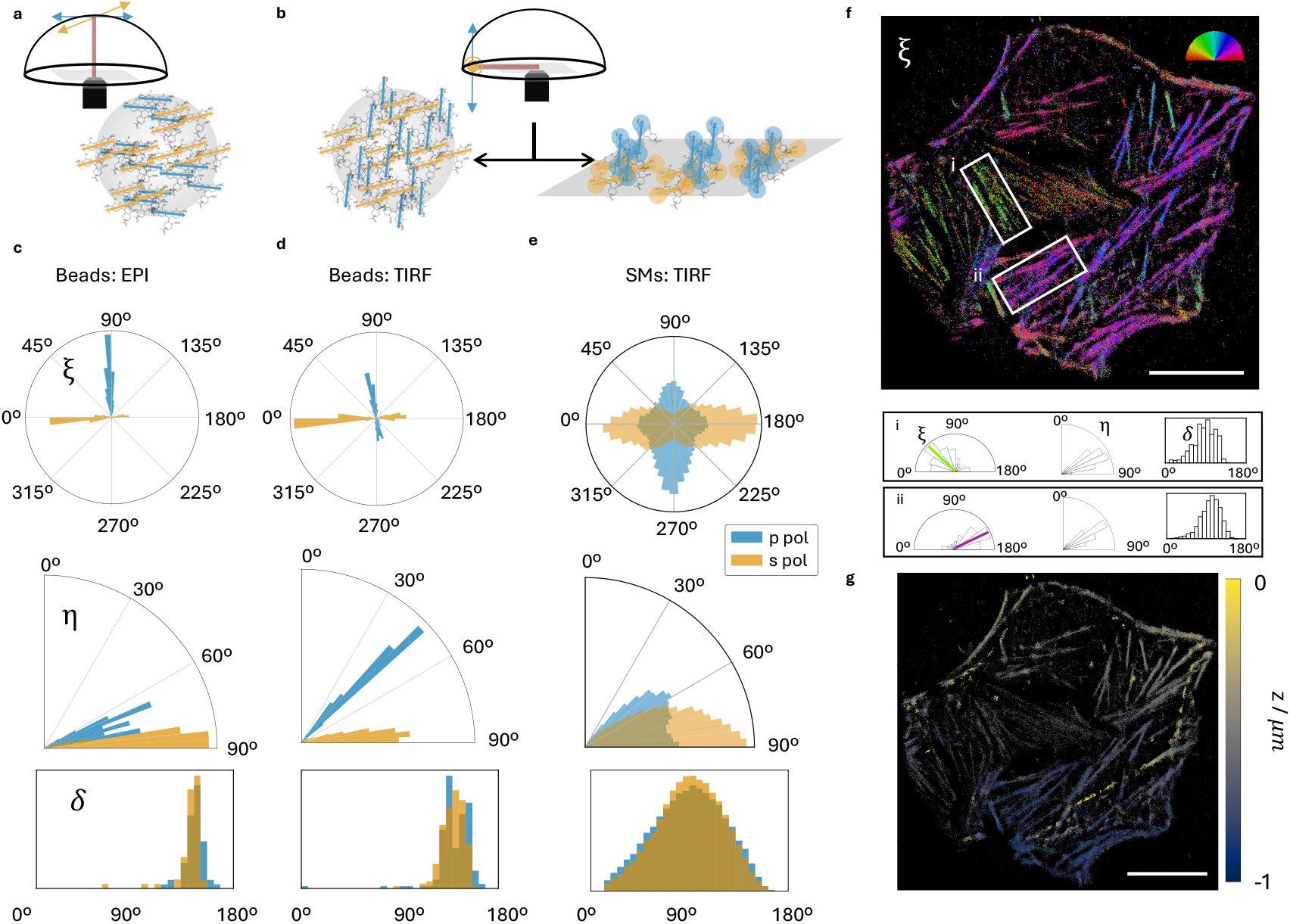
Measurements of polarization-dependent illumination and orientation retrieval. **a** *Epi-illumination schematics*. The optical configuration under normal incidence illumination, with yellow and blue arrows indicating, respectively, the s and p polarizations. The diagram illustrates how these polarizations excite the fluorophores of fluorescent tetraspeck beads differently under epi conditions. **b** *TIRF illumination schematics*. The total internal reflection configuration with the same s (yellow) and p (blue) color code. In TIRF, the p-polarized field is perpendicular to the sample plane, exciting perpendicular fluorophores, while the s-polarized field lies parallel to it. Representative images show the distinct excitation effects for p and s polarizations on fluorescent beads (left) and on single Alexa Fluor 568 molecules (SMs) immobilized on a coverslip (right). **c–e** *Orientation analysis*. Maps of in-plane orientation angle (*ξ*), out-of-plane angle (*η*), and wobble (*δ*) derived from measurements on three systems with s and p polarization schemes described in a-b: beads in epi illumination, beads in TIRF illumination, and single molecules in TIRF illumination. A total of 482 beads and 111,099 single-molecule detections were used in the analysis. **f** *SMOLM-LFM imaging of a fixed U2OS cell labeled with phalloidin–AF568*. In-plane orientation angle (*ξ*) of single emitters is represented by the color code shown in the top-right corner. Insets represent quantitative maps of the orientation parameters, i.e. in-plane (*ξ*), out-of-plane (*η*) angles, and wobbling angle (*δ*), corresponding to the regions highlighted. **g** The same cell as in f, color-coded according to the axial (z) position of the localizations. Scale bars: 20 µm.

In contrast to Monte Carlo simulations, the Stokes basis matrix to be used for the parameter’s retrieval in the experimental situations described here does not correspond exactly to the ideal theoretical matrix. Indeed in experimental measurements, microscope- and sample-induced polarization distortions, as well as BFP non-idealities, introduce deviations that must be accounted for. Because SMOLM-LFM does not use polarization-resolved detection, and therefore cannot rely on standard polarimetric calibration procedures, a specific experimental calibration of the Stokes basis is required to ensure accurate orientation retrieval. In practice, we experimentally estimated the correction to the Stokes basis matrix based on known physical situations, as detailed in Supplementary Note 2.

Figures 3 c–e show the histograms of the retrieved mean orientation parameters. For beads in epi (normal indicence) illumination, as shown in Fig. 3c, the measured in-plane angles *ξ* match well with the expected values. Since the illumination fields are parallel to the sample plane, *η* should ideally be 90°; this estimation is accurate for s polarization and slightly underestimated for p. The measured wobbling angle is *δ* = 140 *±* 13° on average, indicative of strong depolarization which can be explained by both the photoselection angular width and the remaining energy transfer between fluorophores in the beads. Nanobeads under TIRF illumination (Fig. 3d) show the expected behavior, with p polarization preferentially exciting vertically oriented fluorophores whose *ξ* angle is poorly defined. For such emitters, the intensity near the center of the field-of-view is low, making them difficult for the SMLFM algorithm to detect. Consequently, the measured out-of-plane angle *η* = 45 *±* 7° represents an effective limit of the technique in realistic conditions, as vertical fluorophores emit weakly in the central channel. Lastly, randomly oriented single fluorophores on a coverslip were measured under TIRF illumination (see Materials and Methods and Fig. 3e). For s polarization, the retrieved parameters were *η* = 72 *±* 12° and *δ* = 90 *±* 30° which is in agreement with the expected spread arising from the orientation distribution of this type of sample (8). For the p polarization, the retrieved parameters were *η* = 64 *±* 15° and *δ* = 89 *±* 31°. Note that, as observed for the bead measurements, the *ξ* angle is poorly defined for more vertically-oriented fluorophores, resulting in a more homogeneous distribution of *ξ* values. At last, the single molecule’s sample exhibits larger widths of histograms compared to those obtained with nanobeads: this is consistent with the measured SNRs, which are respectively 1.5 and 10.

### Imaging actin inside a cell

To further validate the capability of SMOLM-LFM for complex biological imaging, we applied it to measure the intracellular 3D orientation and spatial organization of actin filaments. For this, acting filaments in fixed U2OS cells were labeled with phalloidin-Alexa Fluorophore 568 (AF568) to measure the local orientation of individual fibers (see Materials and Methods). Figure 3f shows that the in-plane orientation of the filaments is accurately retrieved across a field-of-view (FOV) larger than 50 µm. Insets

(i) and (ii) highlight two regions containing fibers oriented along nearly perpendicular directions, both exhibiting similar average wobbling angles: *δ* = 87 *±* 24° for region (i) and *δ* = 99 *±* 22° for region (ii). Similarly, the in-plane orientation, *ξ*, which follows the local direction of each filament, shows similar values across both regions. The mean out-of-plane angles for the two regions are *η* = 63 *±* 11° for fibers aligned with the direction of the illumination tilt (i), and *η* = 62 *±* 10° for those oriented perpendicularly (ii). Those out of plane angles are consistent with actin filaments almost oriented in the sample plane, as expected from ventral stress fibers in adherent cells. Figure 3g presents the corresponding height map of the reconstructed localizations. The observed tilt in height originates from the deliberate inclination of the coverslip supporting the cell, confirming that the technique successfully retrieves both 3D position and molecular orientation within the sample. More complex imaging experiments are left for future work, as the method remains challenging under low-signal conditions due to the seven-channels splitting inherent to the system.

## Discussion

The main strengths of SMOLM-LFM is to discern fluorophores orientation in 3D without complex polarisation optics and that it is a simple, in-line and robust setup. This differs from several SMOLM techniques whose building and alignment are more cumbersome due to the multiplicity of beamlines (splitting and remixing) along the imaging path (7, 8, 26). Additionally, note that polarization sensitive elements on the detection path, such as Wollaston prisms, waveplates and tailored birefringent masks, can be expensive, not readily available, and prone to misalignments and extra polarization distortions. A second strength, and probably the key feature, is that SMOLM-LFM decouples simply and effectively the in-depth (axial) position from the orientation estimation. As shown in Fig. 1, the former leads to a relative displacement of the PSF of different channels, while the latter leads to a change in the relative intensity ratio between them. Moreover, to measure these changes, they both require only a simple fit and linear decomposition that are not computationally expensive. Note that the intrinsic reduced NA of each individual channel leads to an extended depth of field and hence providing the SMOLM technique with a long working range for orientation estimation and localization, which could be used for single molecule tracking, where simultaneous orientation and position are needed (27–29). See Suplementary

Note 3 for a longer comparison between SMOLM-LFM and other techniques.

A first trade-off of the proposed SMOLM-LFM approach is the seven-channel split. Dividing the signal into seven perspective views necessarily reduces the effective FOV (since all channels must fit onto a single camera) (30), lowers the lateral resolution, and, most critically, limits the applicability of the method to dim fluorophores. Note that single fluorophore detection is photon limited, ranging to a few thousand photons that is then divided into the different channels. One possible path forward is to explore alternative MLA geometries with fewer microlenses. As long as more than or equal to four channels are available (which is the minimum required to estimate orientation and wobble under the isotropic wobbling model) the Stokes decomposition remains solvable, however via a non-linear optimization (See Supplementary Note 1). A second important limitation is the non-trivial calibration procedure. Conventional calibration strategies based on polarizers or polarization-sensitive elements (10, 12, 31) are incompatible with our approach. Although we demonstrate workable calibration protocols (described in Supplementary Note 2), further development is needed to make this step more robust and user-friendly. In particular, the estimation of out-of-plane angles remains imperfect and could benefit from improved calibration schemes or additional constraints.

A third limitation is a fundamental detection bias that contributes to the increased bias and STD observed for out-of-plane dipoles (Fig. 2 and Fig. S6). Dipoles oriented along the optical axis emit zero intensity at the center of the BFP, causing the central channel to appear dark. Because our channels-matching procedure requires the presence of a localization in this central channel, events lacking such a detection are systematically discarded. This behaviour constitutes an inherent limitation (shared by many SMOLM approaches) that disproportionately affects nearly vertical dipoles. Nevertheless, we expect that alternative matching strategies could mitigate this effect in future implementations of SMOLM-LFM. A final trade-off concerns the slight coupling observed between 3D position and 3D orientation, as reflected by the increased error in Fig. 2i. Importantly, this coupling does not arise from the optical design but from the simplified 2D Gaussian fitting model used to localize each PSF. For emitters far from focus, the PSFs broaden and flare in the outer views, causing deviations from a Gaussian profile. While this approximation is imperfect under such conditions, it substantially reduces the computational complexity and remains acceptable for most practical applications, as the resulting errors fall within reasonable and scientifically useful bounds. This behaviour is consistent with other SMLFM implementations, as reported in Ref. (15).

A key outcome of this study is the demonstration that SMLFM systems, which are based on a FLFM arrangement, can also retrieve the orientation of single emitters (thus functioning as SMOLM platforms) without requiring any additional optical components or modifications to the setup. This finding suggests that previously-acquired SMLFM datasets, originally analyzed only for 3D position information, could be reinterpreted to extract orientation data, since such information is inherently encoded in FLFM measurements but had remained unused (provided new calibration, as decribed in Supplementary Note 2 is performed on the original system). This contrasts with a recent FLFM implementation combined with a polarization-sensitive camera (32) for ensemble orientation measurements, where the use of polarizing optics inevitably causes photon losses (an undesirable effect for single-molecule imaging), particularly when combined with the intrinsic multi-channel light-field splitting. Future work may focus on extending the method to live-cell imaging and particle tracking, as well as on simultaneous estimation of position, orientation, and spectral information, in analogy with recently proposed approaches integrating colour and position retrieval (16). Additionally, the reconstruction pipeline could be improved by developing a dedicated algorithm that leverages information from all perspective views, rather than relying primarily on the central FOV as in the current off-the-shelf SMLFM linking procedure, which would reduce the detection bias against out-of-plane dipoles and further enhance robustness.

## Materials and methods

### Optical setup

The SMLFM system was built on an inverted microscope (ECLIPSE Ti2-E, Nikon Instruments). Fluorescence emission was collected using a CFI Apochromat TIRF 60*×* oil-immersion objective (NA 1.49), mounted on a motorized z-drive with 10 mm travel (Nikon Instruments) and encoder-controlled step increments of 0.01 *µ*m, providing a z-positioning precision of 10 nm. The detected light is first cropped at the image plane with a circular apperture and filtered with a BP filter (Chroma, ZET488/561nm). The Fourier lens (f=200 mm, ThorLabs) was placed in a 4f configuration with the tube lens (f=200 mm, Nikon) to relay the back focal plane (BFP). A hexagonal microlens array (f=100 mm, pitch=2.8 mm, custom-made by CAIRN Research) was placed in the BFP. The image was detected on an ORCA-Fusion sCMOS camera (Hamamatsu Photonics K.K., C14440-20UP). To control that the excitation was circularly polarized, a *λ*/2 and a *λ*/4 waveplate (Thorlabs, AHWP05M600 and AQWP05M-600) were used sequentially before entering the microscope. The *λ*/2 plate enables the selection of vertical and horizontal polarization when used alone. When used with the *λ*/4 it allows to rotate the orientation of the inner polarization axis of the system to improve the action of the *λ*/4 to generate a circularly polarised or elliptical input light. Samples were illuminated with laser light at 561 nm with different laser powers (ranging from 50 to 300 mW at the source) and measured with NIS-Elements AR 5.30.05 software and the Nikon Perfect Focus System active. Integration times were typically 100 ms for cells and singlemolecule measurements, and 50 ms for beads. Acquisition stacks ranged from roughly 10 frames for beads to as many as 40,000 frames for cells.

### Sample preparation. Cell lines

U2OS cells were cultured in DMEM (Gibco) supplemented with 42 µM gentamicin (Gibco) and 10% foetal bovine serum (FBS; Gibco). All cells were grown at 37 ºC in a 5% CO_2_ humidified incubator.

### Cell culture, fixation and staining

Cells were seeded on an 8 well-chambered cover glass (Cellvis) with a total of 100,000 cells per well. Cells grew until 60% confluency and were fixed with a 4% paraformaldehyde solution (Electron Microscopy Sciences) and Carbonate-Bicarbonate (CB) buffer for 20 minutes and washed three times with PBS. After, cells were incubated over night with 0.5 µM of AF568Phalloidin (Thermo Scientific PN A12380) in a solution of 0.1% Saponin and 10% BSA in PBS.

### STORM imaging buffer

The final composition of the buffer for 4polar-STORM measurements was 100 mM TrisHCl pH 8, 10% w/v glucose, 5 U/mL pyranose oxidase (POD), 400 U/mL catalase, 50 mM *β*-mercaptoethylamine (*β*-MEA), 1 mM ascorbic acid, 1 mM methyl viologen, and 2 mM cyclooctatetraene (COT). D-(+)-glucose was from Fisher Chemical (G/0500/60). POD was from Sigma (P4234-250UN), bovine liver catalase from Calbiochem/Merck Millipore (219001-5MU), *β*-MEA from Sigma (30070), L-ascorbic acid from Sigma (A7506), methyl viologen from Sigma (856177), and COT from Sigma (138924). Glucose was stored as a 40% w/v solution at 4 °C. POD was dissolved in GOD buffer (24 mM PIPES pH 6.8, 4 mM MgCl2, 2 mM EGTA) to yield 400 U/mL, and an equal volume of glycerol was added to yield a final 200 U/mL in 1:1 glycerol:GOD buffer; aliquots were stored at −20ºC Catalase was dissolved in GOD buffer to yield 10 mg/mL, and an equal volume of glycerol was added to yield a final 5 mg/mL (230 U/*µ*L) of catalase in 1:1 glycerol:GOD buffer; aliquots were stored at −20ºC. *β*-MEA was stored as ≈ 77 mg powder aliquots at 20ºC; right before use, an aliquot was dissolved with the appropriate amount of 360 mM HCl to yield a 1 M *β*-MEA solution. Ascorbic acid was always prepared right before use at 100 mM in water. Methyl viologen was stored as a 500 mM solution in water at 4ºC. COT was prepared at 200 mM in DMSO and aliquots stored at −20ºC. Freshly prepared STORM buffer was typically used on the day of preparation and consumed within 3 hours. To ensure that no particles were in the buffer, it was filtered with 200 nm pore size 4 mm PTFE syringe filters (Whatman Puradisc PN 6784 0402).

### Single AF568 fluorophores deposited on a coverslip

Coverslips were first plasma-cleaned to minimize background fluorescence (optional if high laser power or long exposure is used). A 1 nM solution of phalloidin–AF568 in PBS was then added and allowed to dry for several hours to ensure a high effective degree of physisorption to the coverslip; in our case, drying was accelerated by placing the sample under vacuum for 2 h. Even without complete drying, phalloidin binds efficiently to the glass. Immediately before imaging, a drop of miliQ water was added to rehydrate the sample, which can be imaged for up to 30–60 min before desorption occurs.

## Data analysis

### Image reconstruction

The raw fluorescence images acquired were processed using the Picasso software package to generate SMLM reconstructions. Single molecule localisation and fitting were performed using Picasso Localize with settings optimised individually for each dataset: the box side length for single molecule fitting was set to be larger than three times the typical PSF size (11 pixels in this case), and the minimum net gradient was adjusted (typically between 1000 and 3000) to maximize the number of valid detections while minimizing false positives from background noise. Drift correction was applied using redundant cross-correlation, dividing the datasets into 10 segments.

### SMLFM analysis

Given this initial set of 2D localizations, individual emitters were localized in 3D using custom SMLFM Matlab scripts outlined in (14, 15). Briefly, the most likely subset of 2D localizations in different perspective views corresponding to a unique emitter was identified. Provided that this set of localizations contained more than 5 elements, the 3D location of this emitter was calculated as the least-squares estimate to an optical model relating the axial emitter position to the parallax between perspective views. If the residual light field fit error was below 200 nm, the fit was accepted and the subset of 2D localizations was removed. This procedure was repeated for each individual emitter. System and sample aberrations were corrected by subtracting the residual disparity (calculated for data acquired for all emitters localized during the first 1000 frames) from all 2D localizations prior to calculating the 3D light field fit.

### SMLFM 3D optical calibration

A TetraSpeck bead kit (ThermoFisher Scientific, T14792) mounted with a two-part optical cement (refractive index 1.56) was used for calibration, using 200 nm fluorescent beads. For the 3D calibration, the piezo mounted on the microscope was used to scan the sample axially over 10 µm recording 1 frame at 100 ms exposure per 50 nm increment. The data was reconstructed in 3D and plotted against the known movement of the piezo stage. A linear fit was applied to the calibration curve, the gradient of which was a correction factor subsequently applied to all reconstructed data presented in this work. For the orientation calibration, the Tetraspeck beads were imaged with S and P polarization subsequently. Each measurement comprised recording 10 frames at 50 ms exposure, then averaging the frames prior to analysis.

### Orientation estimation

For the orientation retrieval, only emitters with 6 or 7 localizations on the different FOVs were considered. Every FOV localization had associated an uncertainty and photon counts (proportional to the area under the fitted gaussian). The photon counts were used to compute the intensity at every FOV and then used to find the Stokes parameters associated to each emitter, which were then used to find the orientation parameters, as explained in the supplementary note 1 and in the main text.

## Acknowledgements

This work was supported by the European Union’s Horizon 2020 research and innovation program under the Marie Sklodowska-Curie grant agreement no. 101180631 (https://doi.org/10.3030/101180631) attributed to A.G.V. R.H. acknowledge support from the European Research Council (ERC) under the European Union’s Horizon 2020 research and innovation programme (grant agreement No. 101001332) (to R.H.) and funding from the European Union through the Horizon Europe program (RT-SuperES project with grant agreement 101099654-RTSuperES to R.H.). However, the views and opinions expressed are those of the authors only and do not necessarily reflect those of the European Union. Neither the European Union nor the granting authority can be held responsible for them. This work was also supported by a European Molecular Biology Organization (EMBO) installation grant (EMBO-2020-IG-4734 to R.H.). S.B. and L.A. contribution to the work was supported by by the French National Research Agency under the France 2030 Investment Plan, reference ANR-21-ESRE-0002. H.S.H. acknowledges the support by Fundação para a Ciência e Tecnologia (FCT, Portugal) through the FCT fellowship CEECIND/01480/2021. S.D. acknowledges the support of the Biotechnology and Biological Sciences Research Council (BB/X511092/1, UKRI715) and S.F.L. the Royal Society funding (RGF/EA/181021). This study was also funded by FCT through the MOSTMICRO-ITQB R&D Unit (UIDB/04612/2020, UIDP/04612/2020 to ITQB-NOVA) and LS4FUTURE Associated Laboratory (LA/P/0087/2020 to ITQB-NOVA). Large Language Models were used to assist with stylistic improvements and linguistic corrections in the writing process of the originally submitted manuscript. However, no contributions to the content or to the scientific analysis were made using LLMs. The authors thank Charitra Sree Senthil Kumar for valuable discussions on cell protocols.

## Author contributions

A.G.V. co-conceived the project, designed and implemented the simulations and experiments, and developed the calibration and orientation retrieval methodologies. L.A.C. co-conceived the project, led the theoretical modeling, provided the framework and codes for orientation estimation, and contributed continuous technical input throughout the study. A.G.V. and L.A.C. wrote the manuscript, with contributions and feedback from all authors. S.B. discussed the main ideas and made valuable contributions to the implementation of the technique. For the instrumentation, S.D. trained A.G.V. on the SMLFM system and assisted in setting up the LFM system, while H.H. also assisted with the LFM system setup and contributed to implementing the actin labelling protocol. E.B. provided the original simulation code that served as the basis for the simulations developed in this work. S.F.L. enabled the crucial transfer of SMLFM knowledge. R.H. secured funding and supervised the project, supporting it from its conception through to completion.

## Supplementary Note 1: Orientation estimation via Stokes decomposition

In this section we discuss briefly the theory behind the orientation estimation used in this work, more details on the foundation of this approach can be found in previous publications related to microscopy (5) and polarimetry (24, 25). Fluorescence emission behaves as that of a linear dipole, corresponding to the emission dipole of the fluorophore denoted *µ*. However, since all fluorophores wobble, i.e. their direction oscillates in time, it is better to describe them through the second moment matrix (5), whose elements are of the form Γ_*ij*_ =*< µ*_*i*_*µ*_*j*_ *>*, where *< >* stands for time average. This matrix contains the dipole orientation statistics, meaning its mean orientation, described by two angles, the off-plane angle *η* and the in-plane angle *ξ*, and its wobbling, described in its more general case by three parameters: average wobbling, angle and amount of anisotropy in the wobbling. So any ratiometric, i.e., purely intensity-based, orientation measurement requires at least 6 measurements/channels for these 5 parameters plus the total intensity (number of photons).

As it has been described previously in the literature (24), a useful decomposition of the second moment matrix is in terms of the generalized Stokes-Gell-Mann parameters, *S*_*ij*_, which for a linear dipole is

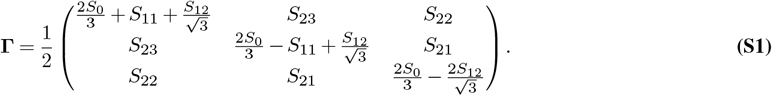

This decomposition offers multiple advantages, since any intensity measurement, **I** = (*I*_1_, …, *I*_*N*_ )^⊤^, where *N* is the number of channels (^⊤^ stands for transpose), can be decomposed linearly in terms of the Stokes basis components, ***I***_***ij***_ = (ℐ _*ij*,0_, …, ℐ_*ij,N*_ )^⊤^, in which the coefficients correspond exactly to the generalized Stokes parameters, i.e.,

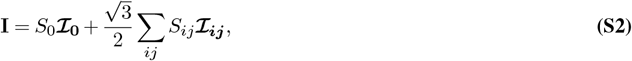

or

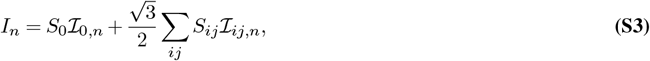

where the subindex *n* indicates the channel/sub-image. To obtain the Stokes basis elements, one needs to compute

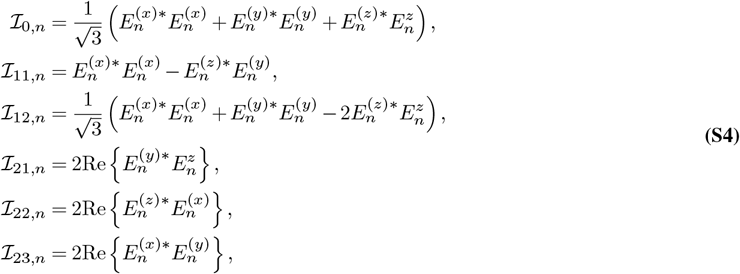

where 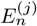 stands for the electric field at the detector coming from a linear dipole oriented along the *j* Cartesian component for a propagating field through the channel *n* of the MLA (^∗^ stands for conjugate). To obtain the electric field at the detector a forward model is used, which has been extensively described in the literature (5, 12, 24).

The first advantage of this decomposition is its computational efficiency: performing a linear decomposition is far less demanding than conventional non-linear fitting routines. The second advantage is conceptual: the Stokes basis elements provide a direct way to analyse and understand the performance and limitations of a given technique. A convenient merit function that can be derived from this linear decomposition is the condition number of the matrix composed of the Stokes basis elements. We used the condition number to compare different microlens arrays (MLA) of same honeycomb geometry but of different sizes (effective numerical apertures), as shown in Fig. S1. Note that this merit function permits to assess the relative effect of experimental parameters on the Stokes parameter’s retrieval efficiency, however to quantify the performance of the method, more complete analysis involving the calculation of the Fisher Information matrix (24) or Monte Carlo analyses (Fig. 2 of the main text and Fig. S4) are required.

With respect estimating the intensity at each channel, *I*_*n*_, either box integration or 2D Gaussian fitting can be used. Depending on the strategy, the way of handling the background differs. For the 2D Gaussian model, the background can be estimated simultaneously and hence signalled out. Note that the 2D Gaussian model is just an approximation, and that it starts failing for very defocused emitters, as shown in Fig. 2 of the main text and Fig. S3. Once the intensities are estimated, we first need to write Eq. S2 in a matricial way:

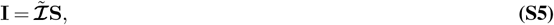

where **S** = (*S, aS, aS, aS, aS, aS* )^⊤^ is the Stokes vector, where 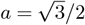, and 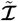 is the matrix made of the Stokes basis elements. Here, since *N* = 7, the pseudo-inverse of 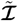 can be computed such that the Stokes vector can be obtained from the intensity measurements as

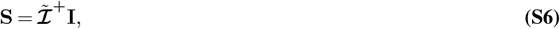

where ^+^ denotes the pseudo-inverse. In this case, we are using only the events in which the 7 channels have been detected and estimated. However, this leaves out many events in which the detection has failed to give a read out in one or multiple channels. In order to increase the number of events analysed, and still benefit from the linear decomposition, this treatment can be immediately extended to those in which only six channels have been detected by

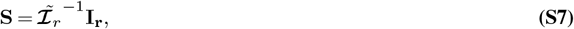

where 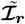 is the matrix made of the Stokes basis elements constructed using only the 6 channels in which a PSF was detected successfully, and **I**_**r**_ analogously but with the intensities. In this case, 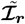 is a 6 × 6 matrix, which can be inverted. For events in which less than 6 channels detections, orientation can still be carried out, however, no longer with a straightforward linear decomposition (refer to the end of this supplementary note for a longer discussion). Note that the quality of the linear decomposition on the Stokes basis can be estimated for each detection by computing the RMS error of the fitted intensities, **I**_est_, with respect the measured ones. We choose to use a minimized RMS version (by a multiplicative factor, *c*, to prioritize looking for distribution similarities), i.e.

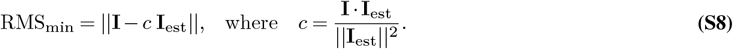

The last step to retrieve the orientation parameters, (*η, ξ, δ*), is to convert the Stokes parameters into the physical orientation angles and wobble. In the widely-used wobbling-inside-a-cone model, in which wobble is only characterized by an oscillation cone angle, *δ*, closed form solutions relating the Stokes parameters and the angles have been described in the literature (24). Here we estimate the angle parameters through the construction of the second moment matrix (using Eq. S1) and from it to compute its eigenvalues, Λ_*i*_, and eigenvectors, **e**_*i*_, since it takes into account all Stokes parameters at once rather than selecting a few to retrieve the angles. In this computation, the eigenvector, **e**_1_ = (*e*_1,*x*_, *e*_1,*y*_, *e*_1,*z*_), with the largest eigenvalue, Λ_1_, corresponds to the mean dipole orientation, described by *η* (out-of-plane angle) and *ξ* (in-plane orientation). Explicitly,

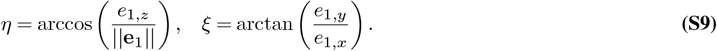

where || || stands for the norm. The two remaining eigenvectors and eigenvalues describe the wobbling (24). In order to simplify the understanding of the wobbling, it is useful to assume the cone model for isotropic wobbling, in which the angle of the cone is given by

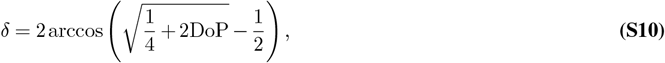

where DoP is the degree of polarization,

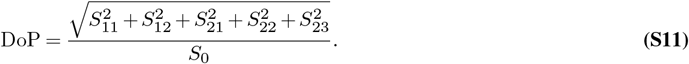

Note that under the assumption of isotropic wobbling (Λ_2_ = Λ_3_ theoretically, although in practice they are never exactly equal), the degree of polarization is equal to

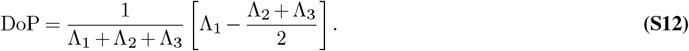

### Minimum number of channels for orientation retrieval

Although the minimum number of intensity channels to fully retrieve the intensity, orientation and wobble of an emitter is six, corresponding to the six real generalized Stokes parameters, under the isotropic wobbling assumption or under the wobbling-in-a-cone model, the number of channels is reduced to only four, corresponding either to (*I, η, ξ*, DoP) or (*I, η, ξ, δ*), respectively. However, as described in the literature (24), the dependence becomes non-linear, thereby requiring a more computationally expensive and time-consuming non-linear optimization.

## Supplementary Note 2: Calibration of the orientation parameters retrieval

Using the theoretical Stokes basis to retrieve molecular orientations is a priori not accurate enough to recover the correct orientations parameters, as illustrated in Figs. S2a, d. In this case, the basis was generated using the nominal parameters of the microlens array (MLA) and objective lens to represent an ideal system. When this ideal Stokes basis is applied to the experimental calibration measurements on fluorescent beads under EPI and TIRF illumination with s and p polarizations (see main text), the retrieval yields incorrect orientation parameters for all conditions, as shown in Fig. S2d.

In practice, the MLA used in this study deviates from the ideal design due to fabrication imperfections. As shown in Fig. S2b, the real array is cropped at the edges, reducing the effective aperture of the microlenses at the back focal plane. Nevertheless, the impact of this geometrical deviation is minor. As illustrated in Fig. S3a, the retrieval errors obtained using the full and cropped pupils exhibit very similar trends across the axial coordinate z for the same photon count, indicating that the influence of the MLA cropping on orientation retrieval performance is negligible.

Incorporating the actual MLA geometry into the forward model improves the retrieval, as shown in Fig. S2e, but some illumination conditions such as the p-polarized EPI excitation still produce deviations from the expected values. A complete calibration must therefore account not only for the correct MLA geometry but also for real optical distorsions introduced by the experimental system. To this end, we performed a calibration procedure to reweight the Stokes basis, compensating for system-specific imperfections. This was achieved by acquiring three independent measurements of Tetraspeck nanobeads with three different illumination schemes corresponding to three orthogonal emitter orientations:

1. epi configuration with linear s polarization in the sample plane,
2. epi configuration with linear p polarization in the sample plane, and
3. TIRF configuration with a p polarization to provoke an equivalent orientation measurement perpendicular to the sample plane.

These calibration measurements are schematized in Fig. S2c. The measured intensities from these three configurations were used to redefine the weighting of the experimental Stokes basis. Once this empirically calibrated basis was introduced, the retrieval of orientation parameters from unknown samples (Fig. S2f) reproduced closer-to-the-expected dipole orientations, demonstrating the necessity and effectiveness of system-specific calibration for reliable orientation estimation.

For the three calibration scenarios (mean *±* STD, *n* = 482), the normalized RMS errors (RMS error of the intensity-ratios fitting using the Stokes-basis decomposition, normalized by the average photon count for each condition) were (1.106 *±* 4.09) *×* 10^−9^ for the experimental calibration, (1.428 *±* 4.80) *×* 10^−9^ for the theoretical cropped MLA, and (3.53 *±* 5.80) *×* 10^−9^ for the theoretical full MLA. These results show that relying solely on the theoretical Stokes basis leads to larger and more variable errors. Previous calibration procedures, which use polarizers or polarization sensitive/transforming masks, cannot be used for SMOLM-LFM: indeed the working principle for the other procedures is based on the properties of polarization, whereas here, the working principle is the intensity distribution at the BFP. Lastly, as we discuss in supplementary note 3, this new calibration methodology are not essentially a drawback, since most SMOLM techniques use samples with known orientation and spatial distribution to refine their calibration (which requires knowledge in the preparation, measurement procedures and optimization algorithms).

## Supplementary Note 3: Comparison tables between 3D SMOLM methodologies

Table S1 presents a qualitative comparison between existing 3D SMOLM techniques (3, 8, 10, 11, 26). Here, we focus on methods capable of estimating the 3D orientation of single emitters while also providing some degree of axial localization. We emphasize that an important bottleneck, both for post-processing and for broad dissemination among users, is the need for dedicated PSF fitting of single-molecule events. Such fitting requires more complex models and therefore (1) slows down data analysis and (2) limits adoption (particularly by non-specialists interested primarily in applying these methods to real biological samples).

**Table S1.**
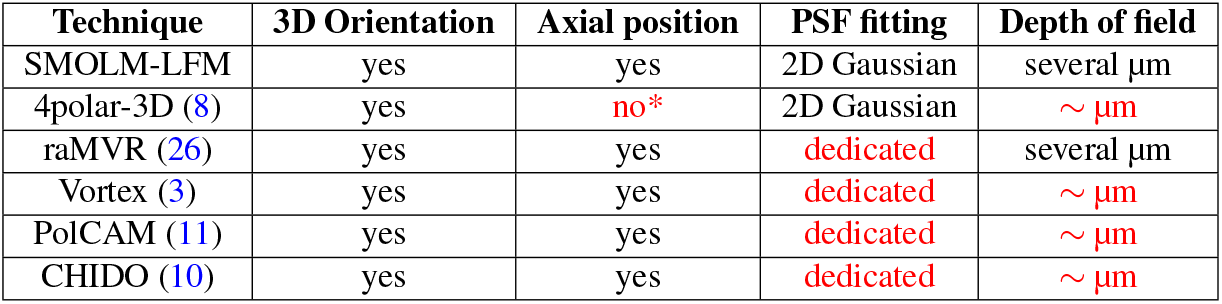
Comparison between optical setups of several different SMOLM techniques: 4polar 3D(8), raMVR(26), Vortex(3), PolCAM (11) and CHIDO (10). Here we compare the capabilities of estimation of each technique, while also highlighting the PSF fitting analysis required per event and the depth of field in which the techniques have been tested. *Not yet fully developed, although it is sensitive.

In this respect, ratiometric techniques such as 4polar-3D (8) and SMOLM-LFM (which require only intensity estimation while preserving a simple PSF shape) enable fast and straightforward fitting, for example using standard 2D Gaussian models. These simple models are already implemented in most SMLM plugins commonly used in the community, making them particularly attractive. Moreover, reducing the complexity of the fitting model lowers computational cost and increases throughput.

Another important aspect is the depth of field (DOF), i.e., the axial range over which the technique maintains performance close to its optimal conditions. Because SMOLM-LFM uses sub-BFP microlenses, the effective NA of each channel is reduced, which in turn increases the DOF.

Beyond data analysis and measurement capabilities, a second set of relevant characteristics relates to setup complexity and required optical elements, summarized in Table S2. Inline geometries are generally preferred because they are easier to build and align, and they remain more robust over time. Conversely, increasing the number of detection channels reduces the field of view and divides the number of detected photons among them, making such approaches less compatible with low-photon regimes typical of single-molecule experiments.

**Table S2.** Comparison between optical setups of several different SMOLM techniques: 4polar 3D(8), raMVR(26), Vortex(3), PolCAM (11) and CHIDO (10). Here we compare the setup building geometry and elements. PBS, polarizing beam splitter; HWP, half-wave plate; QWP, quarter-wave plate; W, Wollaston prism; VaWP, varying-wave plate; VWP, vortwex wave plate; PolCAM, polarization sensitive camera; MLA, microlens array.

| Technique | Setup geometry | Channels | Cameras | Pol. sensitive elem. | Custom made elem. |
| --- | --- | --- | --- | --- | --- |
| SMOLM-LFM | in-line | 7 | 1 | — | 1 MLA |
| 4polar-3D (8) | complex | 4 | 1 | PBS, HWP, W | — |
| raMVR (26) | complex | 8 | 2 | PBS, VaWP, VWP | 2 sector mirrors |
| Vortex (3) | in-line | 1 | 1 | — | vortex plate |
| PolCAM (11) | in-line | 4 | 1 | PolCAM | — |
| CHIDO (10) | in-line | 2 | 1 | QWP, W | birefringent mask |

Finally, the number of dedicated optical components and detectors is a major practical constraint. Additional (often expensive) elements reduce accessibility, and their availability may not always be immediate. Moreover, the more optical components included, the higher the likelihood of alignment errors and the longer the required setup time. In this regard, SMOLM-LFM uses a simple inline optical architecture requiring only a single dedicated component, namely, the microlens array (MLA).

The last point to address concerns the novelty of the calibration methodology presented in this work. Since SMOLM-LFM does not rely on the polarization signature of fluorescence emission but instead exploits the intensity distribution across the BFP channels, standard calibration procedures based on polarization-sensitive detection (10, 12, 31) cannot be applied. Note, however, that other 3D SMOLM techniques rely on target samples with known orientation distributions to refine their calibration, for example, actin filaments (with known dipole orientations along the fibers) (8, 11) or silica beads coated with lipid bilayers (radial orientation) (11, 26). All such measurements require technical expertise in sample preparation and handling, as well as careful data acquisition and calibration optimization to reach optimal performance. Therefore, the calibration procedure proposed here does not introduce a level of difficulty beyond what is already standard in the field.

## Supplementary Figures

**Supplementary Figure S1.**
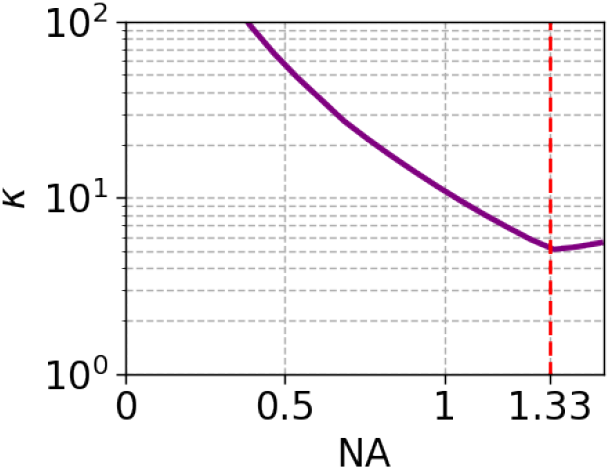
Condition number of the Stokes basis as a function of the NA at the BFP. The total NA corresponds to the effective sum of the NAs of the individual microlenses in the MLA. The region corresponding to the super-critical-angle fluorescence (SAF) is indicated in red. As it can be seen, the best performance is obtained when the total effective NA is around the beginning of SAF.

**Supplementary Figure S2.**
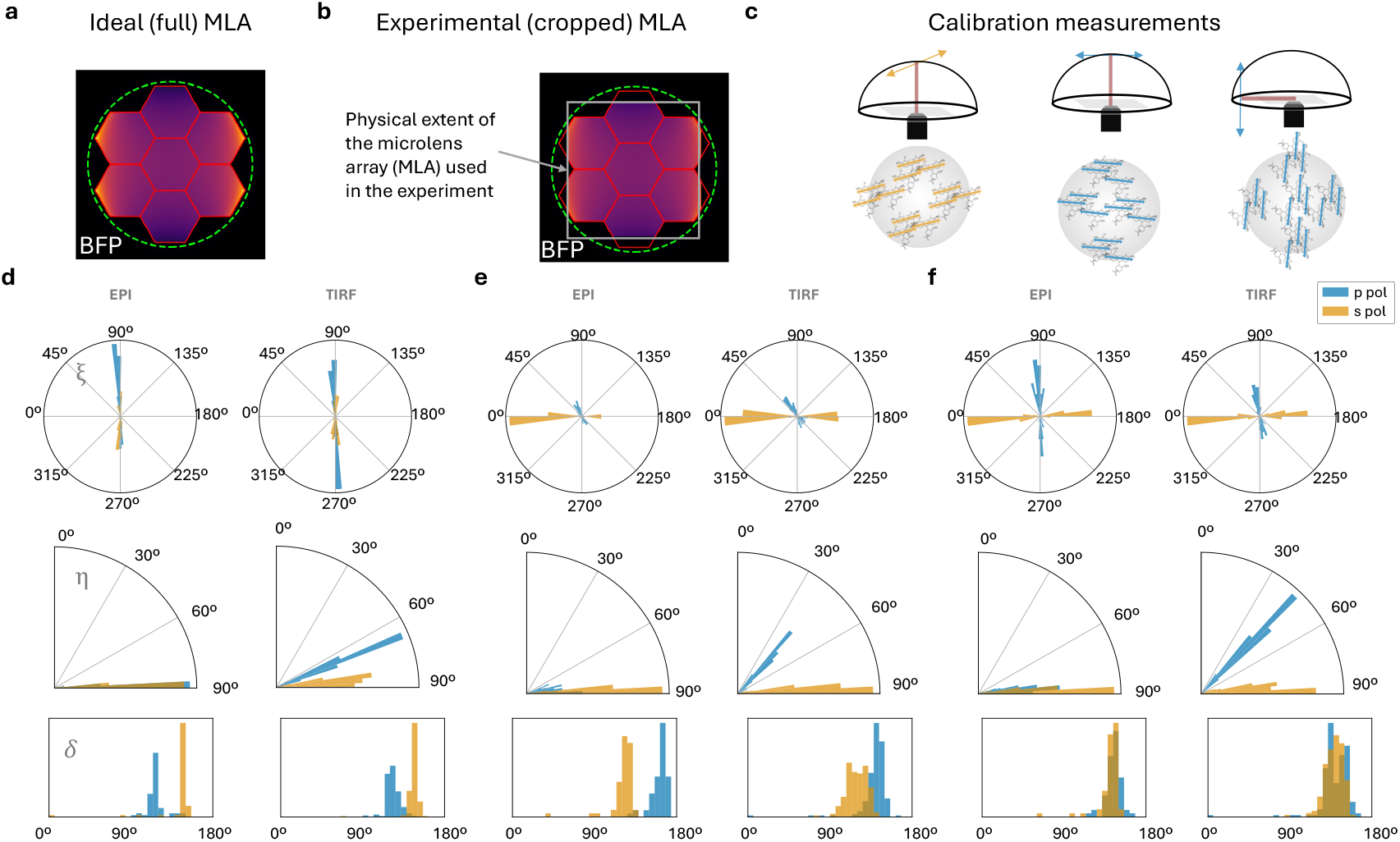
Calibration of the SMOLM-LFM system. Orientation parameter retrieval of photoselected beads with different retrieval information. **a** Intensity distribution at the back focal plane (BFP) of an ideal, complete microlens array (MLA). The green dashed line marks the supercritical angle fluorescence (SAF) region. **b** Intensity pattern at the BFP of the real MLA used in the experiments, showing cropping at the edges due to fabrication imperfections. **c** Calibration measurements on Tetraspeck beads used to retrieve the system response. p and s polarizations in EPI, and p polarization in TIRF, were used to reconstruct the corresponding Stokes parameters of the photoselected emitter populations. **d** Orientation parameters retrieval of tetraspeck beads in EPI and TIRF assuming an ideal, complete MLA. **e** Orientation retrieval using the real MLA shape including the cropped regions. **f** Orientation retrieval using both the real MLA and the experimental calibration data as input, showing improved accuracy in reproducing the measured intensity and polarization distributions. The TIRF histograms were computed from 356 localization events, and the EPI histograms from 126 events.

**Supplementary Figure S3.**
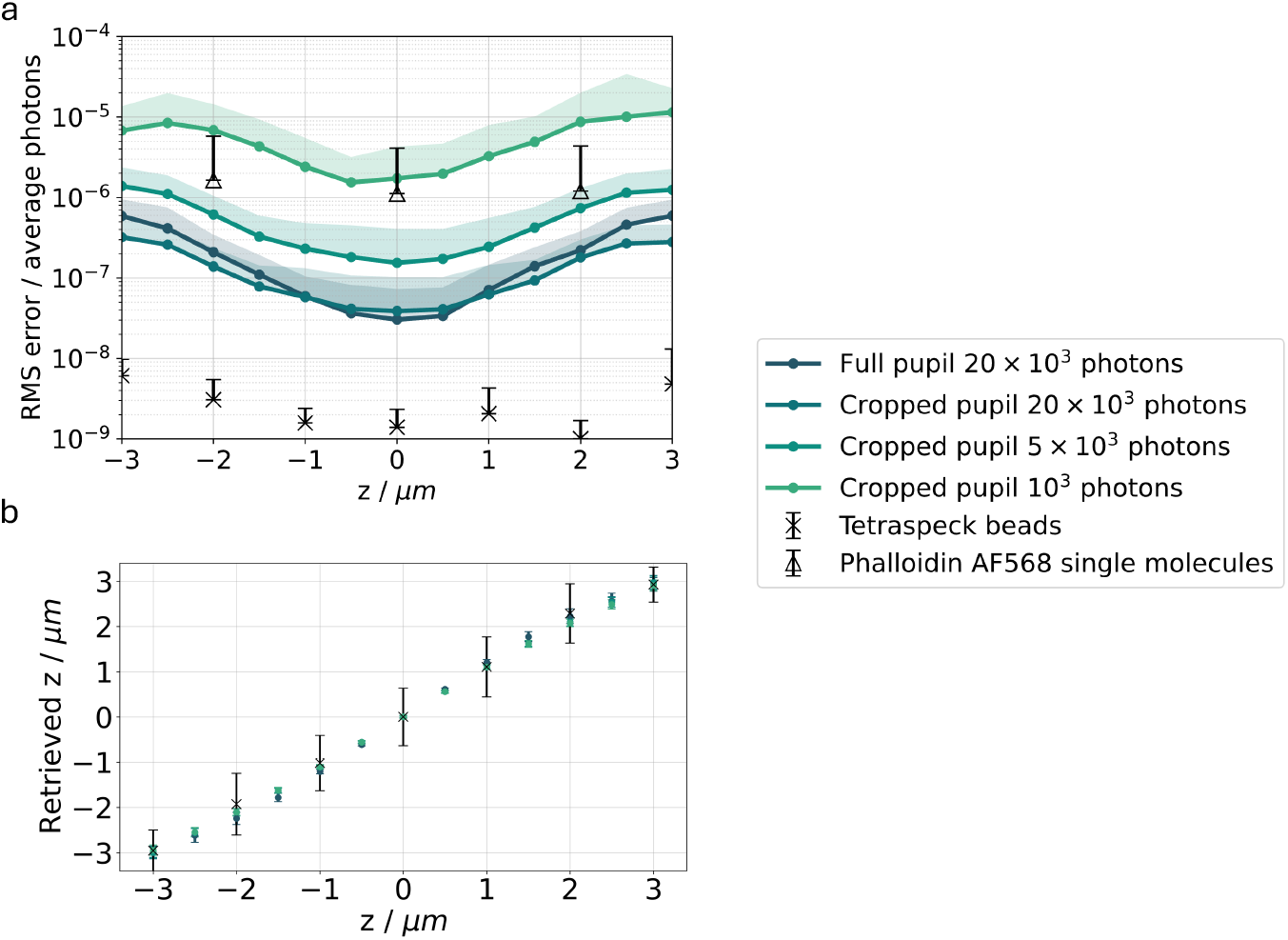
Stokes linear decomposition fitting error across different heights. **a** RMS error of the intensity-ratios fitting using the Stokes-basis decomposition, normalized by the average photon count per condition. Colored dots correspond to Monte Carlo simulations of single dipole emitters imaged and detected across different axial positions (z) for photon counts of 1,000, 5,000, 10,000, and 20,000 corresponding to SNR of approximately 1.5, 2.5, 4, and 7 in the central field of view. Dark blue points represent simulations for an optical system using a full MLA not cropped at the edges, while the other colors correspond to the MLA configuration used in experimental measurements. Crosses indicate experimental data from fluorescent bead measurements (SNR ≈ 10), and triangles correspond to single-molecule measurements of Alexa Fluor 568 (SNR ≈ 1.5). Note that for the single molecule measurements, due to experimental constraints, only three axial positions were used. **b** Retrieved axial position (z) for the Monte Carlo simulations and for measured tetraspeck beads, showing that the z-retrieval accuracy remains consistent between simulated and experimental conditions.

**Supplementary Figure S4.**
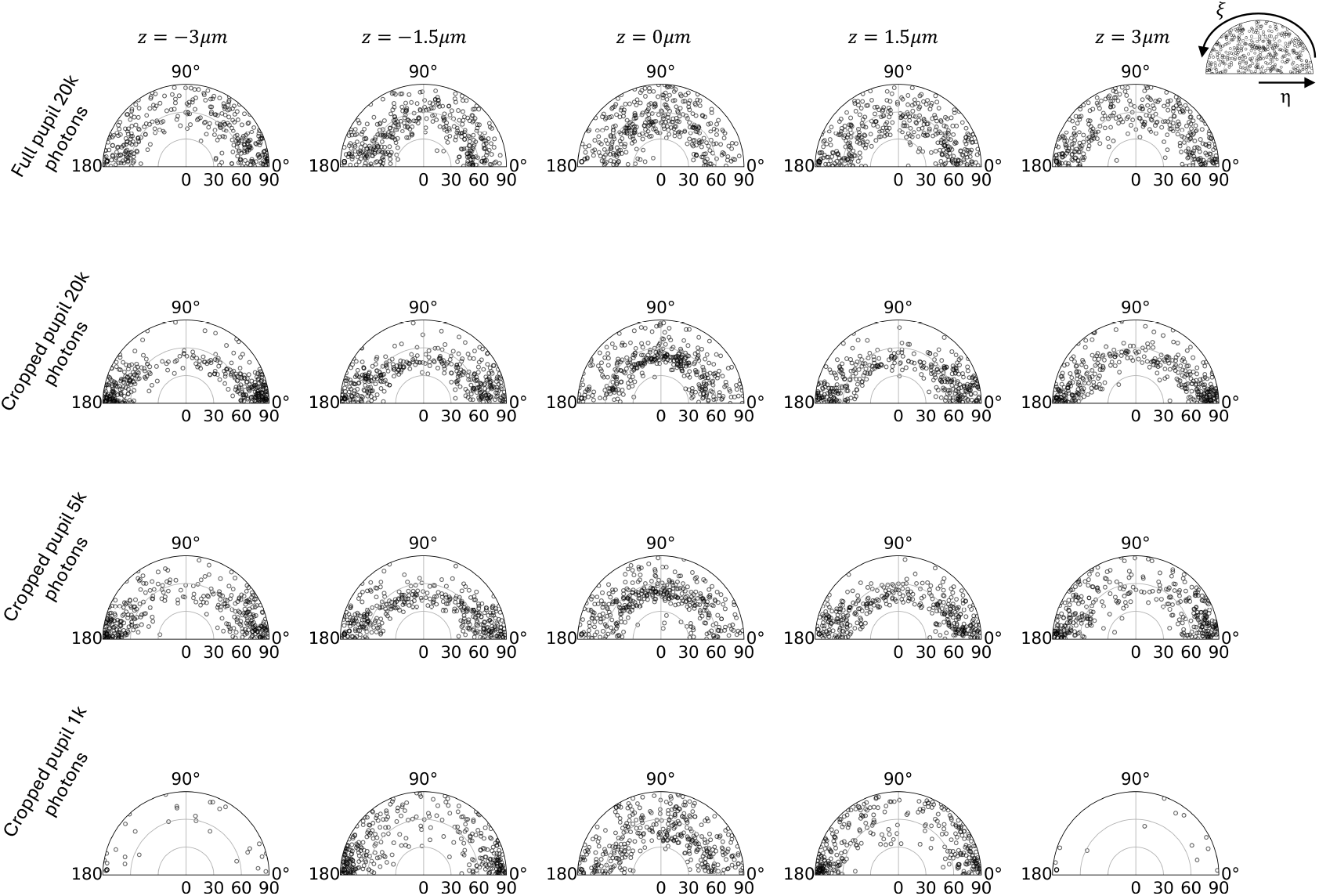
Retrieved orientation parameters across different axial positions. Rows correspond to the simulation conditions using either the full or cropped MLA and photon counts of 1k, 5k, 10k, and 20k, for the same simulations as in Fig. S3. Columns show representative axial positions (*z* = −3, −1.5, 0, 1.5, and 3, *µ*m). Circles indicate the retrieved orientations, with the number of detections ranging from ∼ 1200 at *z* = 0 under the best SNR conditions to progressively fewer points in more demanding regimes. At the focal plane, orientation retrieval remains homogeneous, with the expected limitation that dipoles with *η* ≈ 0^*°*^ are not detected. As defocus increases, the spread of retrieved *ξ* angles becomes less uniform, reflecting reduced sensitivity away from the central plane. Inset (top right): representative angular readout from a homogeneous simulation of 400 emitters, equal to the number of emitters used per condition.

**Supplementary Figure S5.**
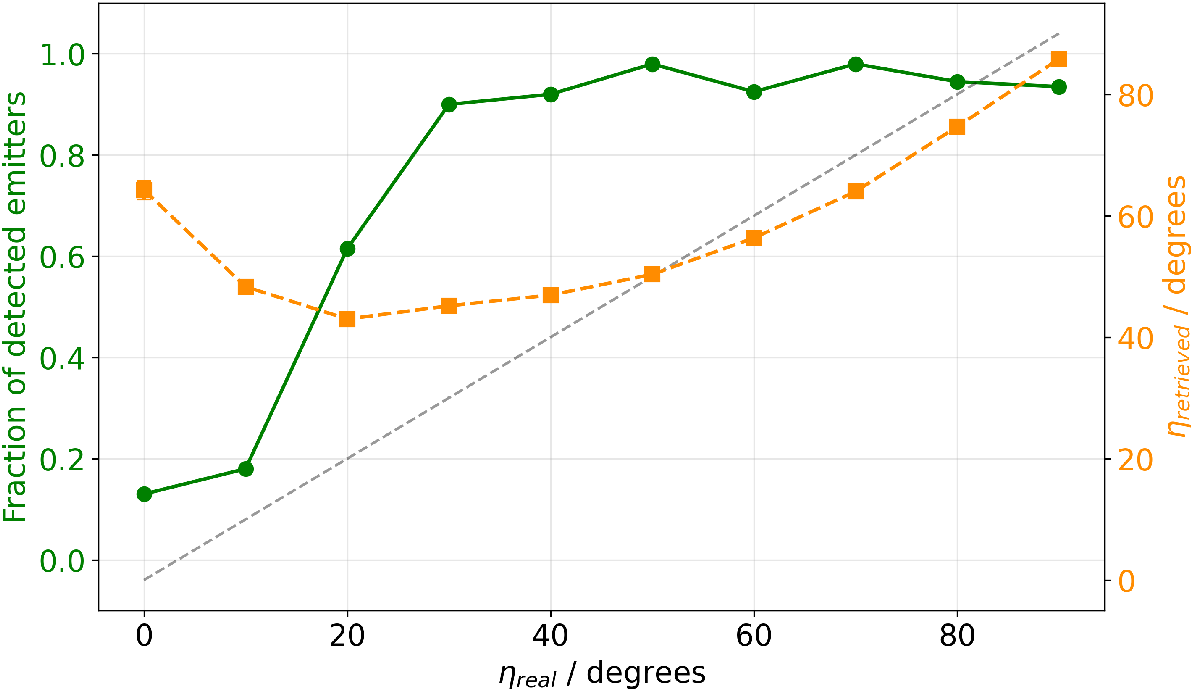
Relation between the number of detections in the central channelcausing the central and the retrieved *η* angle for Monte Carlo simulations mimicking challenging experimental conditions (SNR = 1.8, DoP = 0.7). 200 emitters were simulated per condition. The fraction of emitters producing a detectable localization in the central FOV (therefore yielding a valid multi-view detection set) is shown in green for each simulated orientation. The retrieved out-of-plane angle *η* is shown in orange, with the ideal retrieval indicated by the dashed gray line. These simulations were deliberately carried out in the limiting conditions of single-molecule imaging.

**Supplementary Figure S6.**
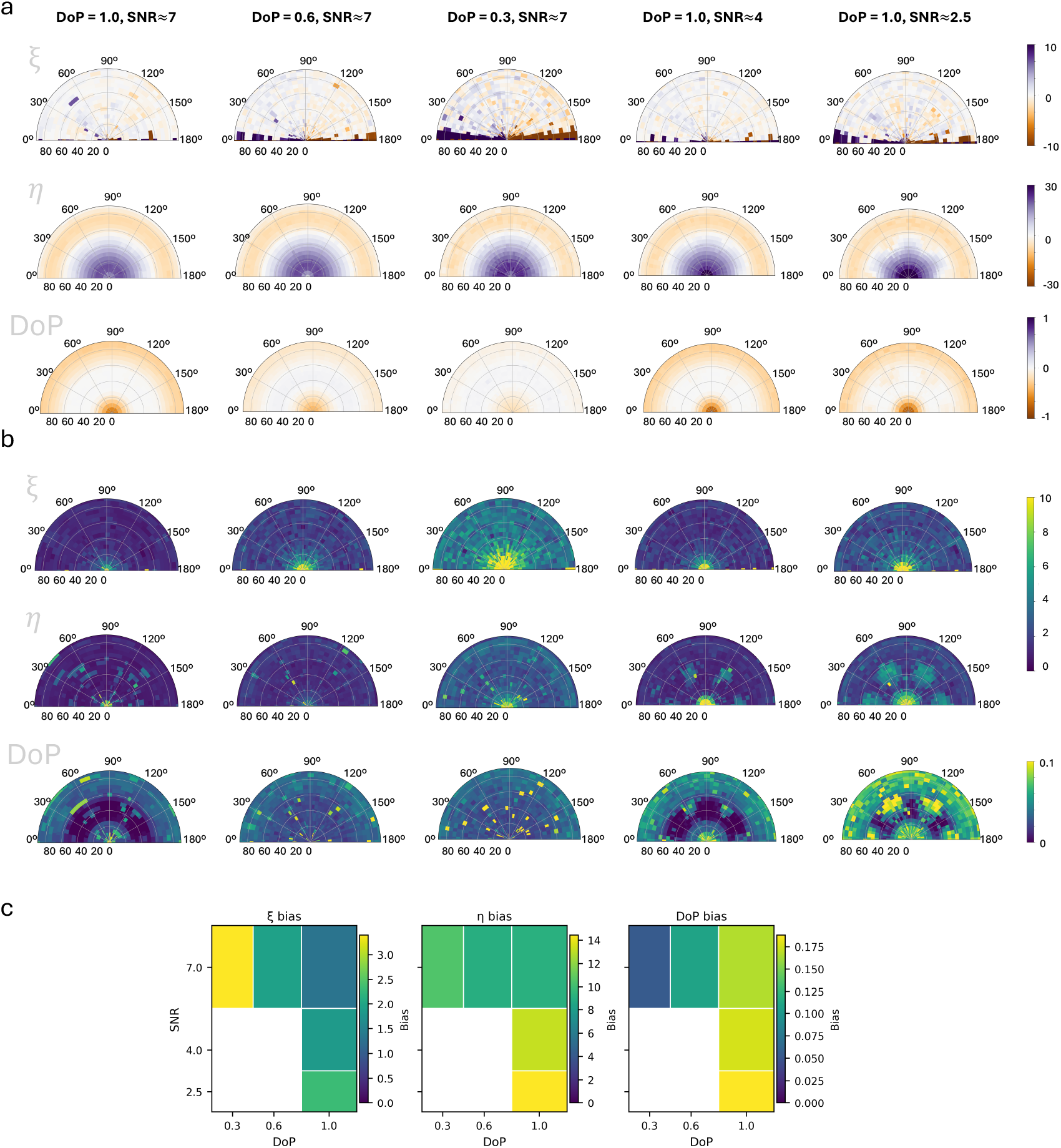
Bias and standard deviation (STD) of Monte Carlo simulations for systems with different SNR and degree of polarization (DoP). **a** Bias of parameter retrieval for emitters detected with 20,000, 10,000, and 5,000 photons, corresponding respectively to SNR of approximately 7, 4, and 2.5. Emitters at SNR ≈ 7 were further simulated for different DoP values. **b** STD of the retrieved orientation parameters for the same conditions. **c** Average bias of the three retrieved parameters (*η, ξ*, and DoP) as a function of DoP and SNR.

